# State-dependent, visually-guided behaviors in the nudibranch *Berghia stephanieae*

**DOI:** 10.1101/2022.10.24.513581

**Authors:** Phoenix D. Quinlan, Paul S. Katz

## Abstract

Nudibranch molluscs have structurally simple eyes whose behavioral roles have not been established. We tested the effects of visual stimuli on the behavior of the nudibranch *Berghia stephanieae* under different food and hunger conditions. In an arena that was half shaded, animals spent most of their time in the dark, where they also decreased their speed and made more changes in heading. These behavioral differences between the light and dark were less evident in uniformly illuminated or darkened arenas, suggesting that they were not caused by the level of illumination. *Berghia* responded to distant visual targets; animals approached a stripe that was at least 15° wide and 50% darker than the background. They did not approach a stripe that was lighter than the background but approached a stripe that was isoluminant with the background, suggesting the detection of spatial information. Animals travelled in convoluted paths in a featureless arena but straightened their paths when a visual target was present even if they did not approach it, suggesting that visual cues were used for navigation. Individuals were less responsive to visual stimuli when food-deprived or in the presence of a food odor. Furthermore, when given a food odor, they had a weaker preference for the dark and behaved similarly in the light and dark. Thus, *Berghia* exhibits visually-guided behaviors that are influenced by odors and hunger state.

**Summary statement:** Behavioral analyses demonstrate that the nudibranch *Berghia stephanieae* is capable of spatial vision and has visually-guided behaviors that are influenced by olfactory information and hunger state.

## Introduction

Gastropod molluscs have been shown to have visual responses ranging from phototaxis (Matsuo et al., 2014; Zieger et al., 2009) to high-resolution spatial vision (Irwin et al., 2021; Land, 1982). Moreover, gastropod species display a wide diversity of eye types ranging from open pit eyes to simple and complex lens eyes (Serb & Eernisse, 2008; Zieger & Meyer-Rochow, 2008). Nudibranchs have relatively simple lens eyes, whose behavioral functions are not known. Studying the role of visually-guided behaviors in nudibranchs has been challenging because animals are often wild-caught, limiting control over the animal’s life history and internal state. To better understand the role of nudibranch eyes, we examined visually-guided behaviors of a laboratory-raised aeolid nudibranch, *Berghia stephanieae* (Valdés, 2005).

Nudibranch eyes are located beneath the integument near the brain. Many nudibranchs lack epithelial pigment over the eye, allowing it to be visible as a small black spot (Hughes, 1970) (Fig. 1). Each eye contains a spherical lens that is covered by a cellular cornea (Chase, 1974; Eakin et al., 1967; Hughes, 1970). Several pigment-producing cells shield light from entering the eye from behind. The eyes of adult nudibranchs possess only three to five photoreceptor cells (Chase, 1974; Eakin et al., 1967; Hughes, 1970), which is fewer than other gastropods that can have hundreds or thousands of photoreceptor cells forming an organized retina (Bobkova et al., 2004; Jacklet, 1969; Meyer-Rochow & Bobkova, 2001; Zhukov et al., 2002). Nonetheless, the positioning and neural connectivity of the photoreceptors in the aeolid nudibranch *Hermissenda crassicornis* suggest that they could support spatial vision (Stensaas et al., 1969; Tabata & Alkon, 1982).

**Figure 1.**
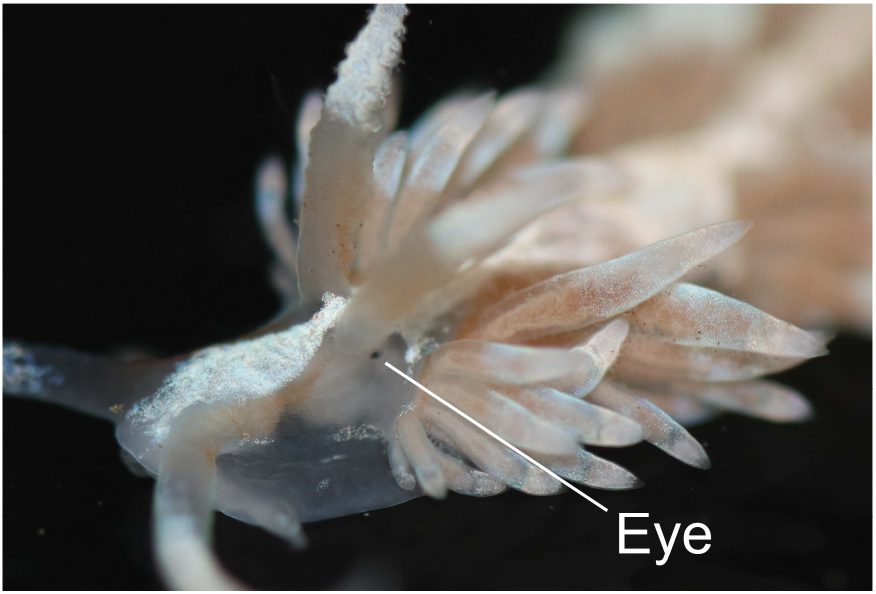
Photograph of adult *Berghia* with eye visible. The eye of adult *Berghia stephanieae* is located dorsolaterally on the head in a non-pigmented zone.

Although spatial vision has not been demonstrated in nudibranchs, they have been shown to have phototactic responses to light. For example, the dorid *Onchidoris bilamellata* spends more time in the dark when given a choice between light and dark (Barbeau et al., 2004), whereas another dorid, *Chromodoris zebra,* and *Hermissenda* spend more time in illuminated areas and approach light sources (Crozier & Arey, 1919; Lederhendler et al., 1980). Furthermore, when *Hermissenda* encounters a shadow in an otherwise illuminated environment, it stops moving forward and returns to the light (Lederhendler et al., 1980).

Anecdotal observations of *Berghia* suggest that it spends most of its time in dark environments, such as underneath objects or in dark crevices. *Berghia’s* responses to visual stimuli have not been experimentally tested. Here, we tested the responses of *Berghia* to visual stimuli to gain insights into the visual behaviors and capabilities of these nudibranchs. We found that *Berghia* exhibits visually-guided behaviors and provide evidence of low-resolution spatial vision. Furthermore, we tested animals under different conditions and found that visually-guided behaviors are state- and context-dependent.

## Materials and methods

### Animal care and husbandry

Specimens of *Berghia stephanieae* were initially obtained from Salty Underground (Crestwood, MO) and Reeftown (Boynton Beach, FL). They were propagated in the laboratory by placing an egg mass into a plastic petri dish and incubating at 30°C. Artificial seawater (ASW) (Instant Ocean, Blacksburg, VA, USA) was maintained at a specific gravity of 1.020-1.022, temperature of 22-26°C, and pH of 8.0-8.5. ASW was exchanged twice weekly through manual pipetting. Late stage juvenile *Berghia* were transferred in groups of ten to 1-gallon acrylic aquariums filled with ASW and kept on a 12:12 LD cycle. *Exaiptasia diaphana* (Carolina Biological Supply, Burlington NC) were added twice weekly as a food source. *Exaiptasia* were kept in glass aquariums filled with artificial seawater maintained at the above conditions. *Exaiptasia* were fed brine shrimp *(Artemia nauplii,* Carolina Biological Supply Co) twice per week.

### Behavioral assays

Individual *Berghia* were video recorded while freely moving inside a circular arena, which consisted of a 9.5-cm diameter glass dish filled with 240 mL ASW. The dish was placed in the center of a piece of 11.5-cm diameter white PVC pipe with a height of 9.5 cm. White cardstock paper was inserted between the pipe and the glass dish. Visual stimuli were printed onto the paper using a Color Laser Jet Pro M454dw (HP, Palo Alto, CA).

An LED tracing board (tiktecklab) was fixed 15.25 cm above the testing apparatus to illuminate the arena. To shade half of the arena, black cardstock paper was placed on top and on one side of the arena. The hemisphere that was shaded was rotated between trials. For tests in illuminated and darkened arenas without a visual target, an 850 nm infrared light (CMVision) illuminated the arena from 30 cm above at a 60° angle. A USB infrared sensing camera with OV2710 CMOS sensor (webcamera_usb) was fixed 16 cm below the dish, recording at 30 frames per second (for trials in a half-shaded arena and uniformly illuminated/darkened arenas) or 2 frames per second (for trials with a visual target).

All animals used were reproductive adults (1-2 cm length) and were tested at least 12 weeks post-hatching. Like other nudibranchs, *Berghia* is hermaphroditic. Each animal was used only once, except when paired testing was performed as indicated. Animals were tested 24-48 hours after being fed. For experiments on food-deprived animals, testing was performed 5-6 days after their last feeding. To create conditioned ASW for food odor, six *Exaiptasia* were kept in 200 mL ASW for 24 hours. 10 mL of conditioned ASW was diluted with 230 mL ASW to provide food odor.

For each trial, a single *Berghia* was gently pipetted to the center of the glass dish. Animals were given 5 minutes to acclimate to the arena, after which they were recorded for 10 minutes. Sample size was chosen using the resource equation approach, which suggested 11-21 animals for within-subjects repeated measures. In the half-shaded arena, 15 animals were tested for each feeding and odor condition (60 animals total). When animals were tested in arenas that were completely illuminated or darkened, the order of the light and dark trial were counterbalanced, and 15 animals were tested.

When testing animals with a visual target, 18 animals were tested for each stimulus type, but individuals were excluded if they did not right themselves immediately upon being pipetted into the arena. No acclimation period was used, and animals were recorded until they reached the edge of the arena or until 6 minutes elapsed. Animals that did not reach the edge were tracked and plotted but excluded from further analysis.

### Analyses and statistics

The location of each individual within the arena was tracked for the duration of the trial using the markerless pose estimation software DeepLabCut (Nath et al., 2019). Networks were trained to detect *Berghia* using training datasets in which animals were manually marked posterior to the first ceratal row. A different network and training dataset were used for each behavioral assay. The trajectories of each animal were exported into CSV files, after which they were analyzed using custom MATLAB scripts. Incorrectly labeled points were removed using criteria such as a likelihood score and the maximum possible distance to travel between frames. For trials with a visual target recorded at 2 frames/second, only *Berghia* that could be tracked from the center to the wall of the arena were included. For trials in a half-shaded arena and uniformly illuminated/darkened arenas, trials were only used if at least 50% of frames (a total of 15000 frames were recorded at 30 frames per second) were labelled correctly. Tracking accuracy was around 98% for fed animals and 88% for food-deprived animals (Fig. S1). Tracking accuracy was higher for fed animals presumably because *Berghia* becomes darker in color after eating, making it more visible against the illuminated background of the video. Arena boundaries were determined by manual segmentation using Make Sense (Skalski, 2019).

To determine whether *Berghia* approached visual targets, the distribution of the locations where *Berghia* touched the wall of the dish were analyzed. Videos were trimmed from when the animal righted itself to when it came in contact with the wall. The trajectories of animals were adjusted so that the first coordinate of each trace was located at the origin, and the location where each animal travelled 95% of the distance to the edge was identified. The R package CircMLE (Fitak & Johnsen, 2017) was used to rank how well 10 models of animal orientation (Schnute & Groot, 1992) describe the distribution of these locations (Table S2). The AICc criterion (Hurvich & Tsai, 1991) was used to compare models, and the model with the highest AICc value was reported as the best fit for the data. Further models were also reported if the relative differences to the best model (ΔAICc) were less than 2, as these models were also strongly supported (Burnham et al., 2011). A visual target was considered to be approached by *Berghia* if the best fitting model was of unimodal distribution directed toward the target.

Behavioral measures such as mean speed, straightness, and mean change in heading were calculated. Mean speed was calculated by dividing total distance travelled by time elapsed. Straightness (straightness index) was calculated by dividing the distance from the center of the arena to the wall by the total distance travelled from the center to the wall. Mean change in heading was calculated as the mean change in direction of 2 vectors defined by the animal’s location across 3 subsequent points in time. Statistical significance was assessed using the Student’s one-sample, two-sample, and paired sample *t*-tests as indicated, with a = .05 (function ‘ttest’ and ‘ttest2’ in MATLAB). A one-way ANOVA was used to assess whether means from multiple groups of animals were significantly different, with a = .05 (function ‘anova1’ in MATLAB). A two-way ANOVA (function ‘anovan’ in MATLAB) was performed to determine the main effect of food-deprivation, the presence of a food odor, and the interaction effect of these two conditions on behavioral measures, including mean speed and mean change in heading. Following the one- or two-way ANOVA, pairwise comparison was performed using Fisher’s Least Significant Difference Test (function ‘multcompare’ in MATLAB) and statistically significant differences were reported. For comparisons of non-parametric data, the Kruskal-Wallis test (function ‘kruskalwallis’ in MATLAB) was used. For detailed results from statistical testing, see supplementary information (Tables S1-S8).

## Results

### *Berghia* preferred dark environments

When placed in an environment that was half-shaded, animals spent most of their time in the dark. The movements of fifteen animals were tracked for ten minutes in an arena that was half-illuminated and half-shaded (Fig. 2A,B). Following a 5-minute acclimation period, 13 of 15 animals (86.7%) were located in the dark side of the arena. During the 10-minute trial, all animals spent a majority of their time in the dark (Fig. 2C). On average, animals spent 83.6 ± 14.2% of their time in the dark half of the arena (*n*=15, one-sample *t*-test: *P*<.001).

**Figure 2.**
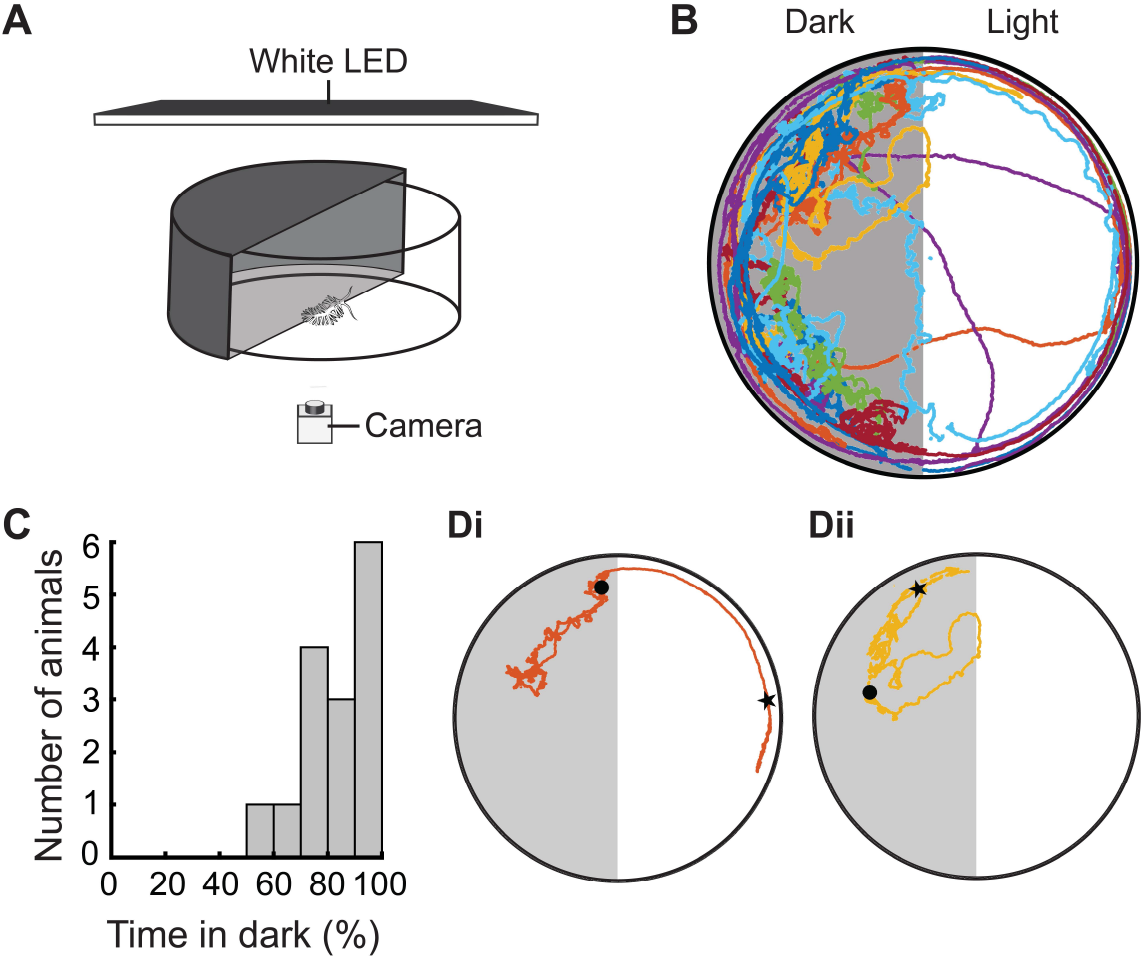
Dark preference in a half-shaded arena. **A.** Diagram of half-shaded arena. The arena was illuminated overhead using white LEDs. A camera was mounted below to record each animal for 10 minutes. **B.** Trajectories of individual animals overlaid, with the darkened (left) and lighted (right) sides marked. **C.** Histogram of the percentage of time spent in dark over 10 minutes (*n*=15). **D.** Example traces from two individuals, one that travelled along the edge of the arena in the light but moved away from the edge and increased turning in the dark (***i***) and one that entered the lighted side, promptly turned around, and re-entered the dark side (***ii***). The starting position (star) and ending position (circle) of each individual is marked.

In the dark half of the area, animals turned frequently and did not stay on the edge (Fig. 2B). In contrast, when animals were in the illuminated half of the arena, their paths were straighter, and they tended to stay near the edge (2B). Fig. 2Di shows an example of an individual that started in the lighted half of the arena, moving along the edge, but once it reached the darkened side, it moved away from the edge and increased the frequency of turns (Fig. 2Di). Fig. 2Dii shows a different individual that started on the dark side, but, after entering the light side, promptly turned around and re-entered the dark. Thus, *Berghia* had a strong preference for being in the dark and showed notable differences in behavior between the light and dark sides.

### *Berghia* behaved differently in uniformly and partially illuminated arenas

To test whether differences in *Berghia*’s behavior could be attributed to the level of ambient lighting, the movements of fifteen animals were tracked in an arena that was either fully darkened or fully illuminated (Fig. 3A). The trajectories of individual animals were more consistent in the dark and light than when in a half-illuminated arena. For example, an individual that circled the perimeter of the arena did so under both the dark and the light conditions (Fig. 3B*i*), and an individual that entered the interior of the arena did so in both conditions (Fig. 3B*ii*). However, in the completely darkened arena, the animals rarely came as close to the edge as they did under uniformly illuminated conditions as can be seen in the individual trajectories (Fig. 3B) as well as the density plots (Fig. 3C*i*).

**Figure 3.**
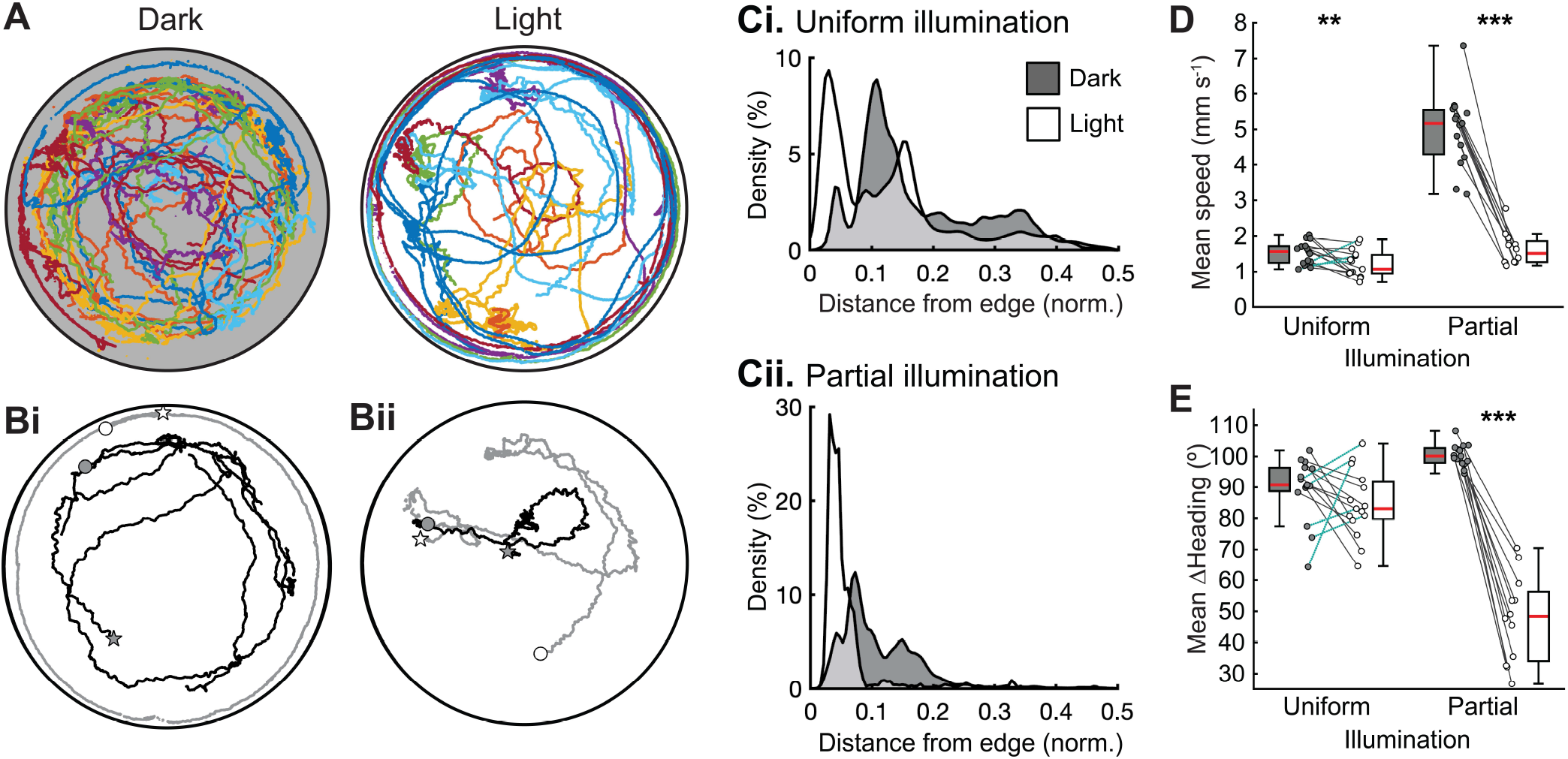
Behavior in arenas with uniform vs partial illumination. **A.** Trajectories of animals (*n*=15) crawling in a completely darkened arena (left) and in one that was uniformly illuminated (right). **B.** Examples of individuals that behaved consistently under both conditions. One individual moved around the edge of the arena in both the light (gray trace) and the dark (black trace), although it got closer to the wall in the light (***i***). A different individual explored the center of the arena under both conditions (***ii***). The start and end positions of each trace are indicated by the star and circle, respectively. **C.** Density plots showing the relative amount of time spent at different distances from the edge of the arena in the light (white) and dark (gray) in an arena that was uniformly illuminated (***i***) and one that was only partially illuminated (***ii***). **D-E.** Box and scatter plots of the mean crawling speed (**D**) and mean change in heading (**E**) of animals in the dark (gray box) and light (white box) in a uniformly versus partially illuminated arena. For all box plots, the median value (red line) is reported. Connected data points are from the same individual; line styles indicate whether the value was higher in the dark (solid gray line) or light (dotted teal line) for each individual. A paired *t*-test was used to test whether mean speed (**D**) or mean change in heading (**E**) was significantly different in the dark compared to the light; significant differences: \*\**P*<.01, \*\*\**P*<.001.

Under uniform illumination, animals frequently made contact with the edge of the dish and spent most of their time within a body’s length (about 1 cm) of the edge of the 9.5 cm-diameter dish (Fig. 3C*i*). However, the proportion of time spent within a body’s length of the edge in the lighted side of a partially illuminated arena was much higher than in a uniformly illuminated one (Fig. *3Ci,ii*).

There were other behavioral differences between animals in uniformly and partially-illuminated arenas. Although animals crawled about 75% faster in a fully darkened arena compared to one that was fully illuminated, the difference was more pronounced in a partially illuminated arena where animals crawled 300% faster in the dark side than the light side (Fig. 3D). Additionally, although animals did not exhibit significantly different turning behavior in uniformly darkened and uniformly illuminated arenas, there was a strong decrease in the mean change in heading when animals were on the light side of a partially illuminated arena, indicating that they turned less in the light (Fig. 3E). Thus, *Berghia* behaved differently in partially-illuminated versus uniformly-illuminated arenas, suggesting that they may be responding to visual features of the environment and not just ambient light levels.

### *Berghia* approached visual targets

Animals were placed in a uniformly illuminated environment with or without a single vertical stripe on the wall outside of the arena (Fig. 4A). Animals placed in the center of an arena with no external markings typically changed directions several times before approaching the edge and 29.4% of them did not reach the wall at all (Fig. 4B). However, with a black stripe that extended 45° around the arena, every animal reached the wall and most animals approached the wall near the stripe, either moving directly toward it or making a large orienting turn before moving in a straight path toward the stripe (Fig. 4C). When tested with a white stripe on a black background, all animals approached the black part of the wall rather than the stripe (Fig. 4D).

**Figure 4.**
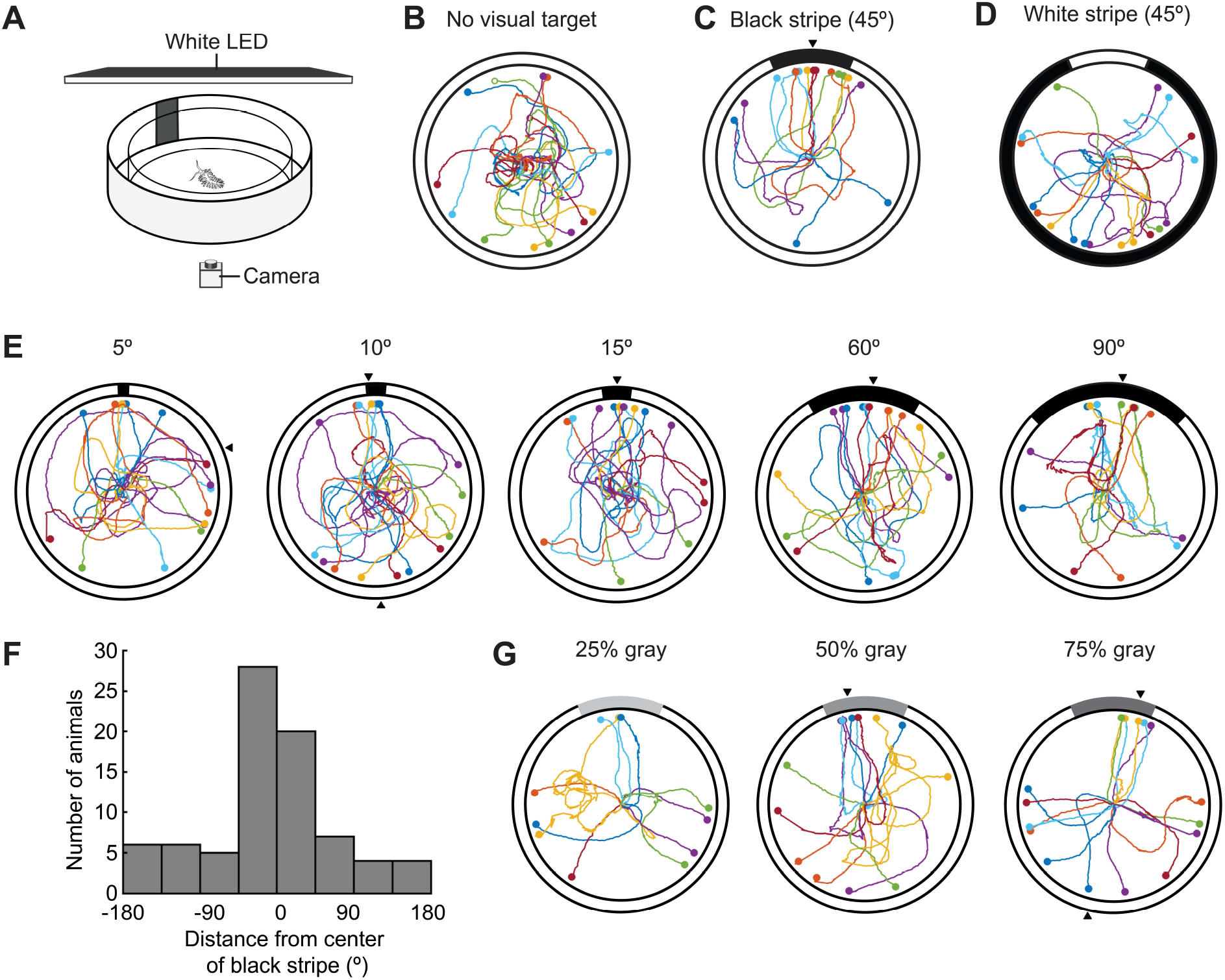
*Berghia* approached visual targets. **A.** Diagram of visual target assay. Animals were placed in the center of a brightly illuminated circular arena surrounded by a white wall with a single vertical stripe. A camera was mounted below to record animals until they reached the wall. **B-D**. Trajectories of animals crawling from the center to the edge of an arena with no visual target (**B**), a 45° black stripe on a white background (**C**), or a 45° white stripe on a black background (**D**). The location where each animal approached the edge is marked with a circle. Animals that did not reach the edge within the allotted time were plotted (open circles) but excluded from analysis. **E.** Trajectories of animals crawling from the center to the edge of an arena with black stripes of different widths. **F**. Histogram of the locations where 97 animals approached the edge of an arena with a black stripe with a width of 15-90°. **G**. Trajectories of animals crawling in response to 45° stripes of different levels of gray on a white background. Maximum likelihood analysis of circular data was used (**B-D,F,G**; Table S2); black triangles mark the direction(s) of best-fitting model for unimodal and bimodal models. Sample sizes were *n*=17 for none; *n*=17 for 45° black; *n*=18 for white; *n*=17, 17, 15, 18, and 14 for 5°, 10°, 15°, 60°, and 90° black, respectively; *n*=12, 16, 17 for 25%, 50%, and 75% gray, respectively.

Animals were tested with stripes of various widths and contrast from the background (Fig. 4E). Animals approached a black stripe that was at least 15° of the circumference of the arena. While some animals approached a 10° stripe, most animals went to the opposite side, and the locations where animals approached the edge followed an axial distribution. Animals approached the widest stripe tested, which was 90°. The locations where animals reached the wall followed a unimodal distribution that was centered on the visual target (Fig. 4E). Animals did not approach a 25% gray stripe but approached a stripe that was 50% gray or darker (Fig. 4G). Overall, *Berghia* most reliably approached a 45° black stripe on a white background (Fig. 5).

**Figure 5.**
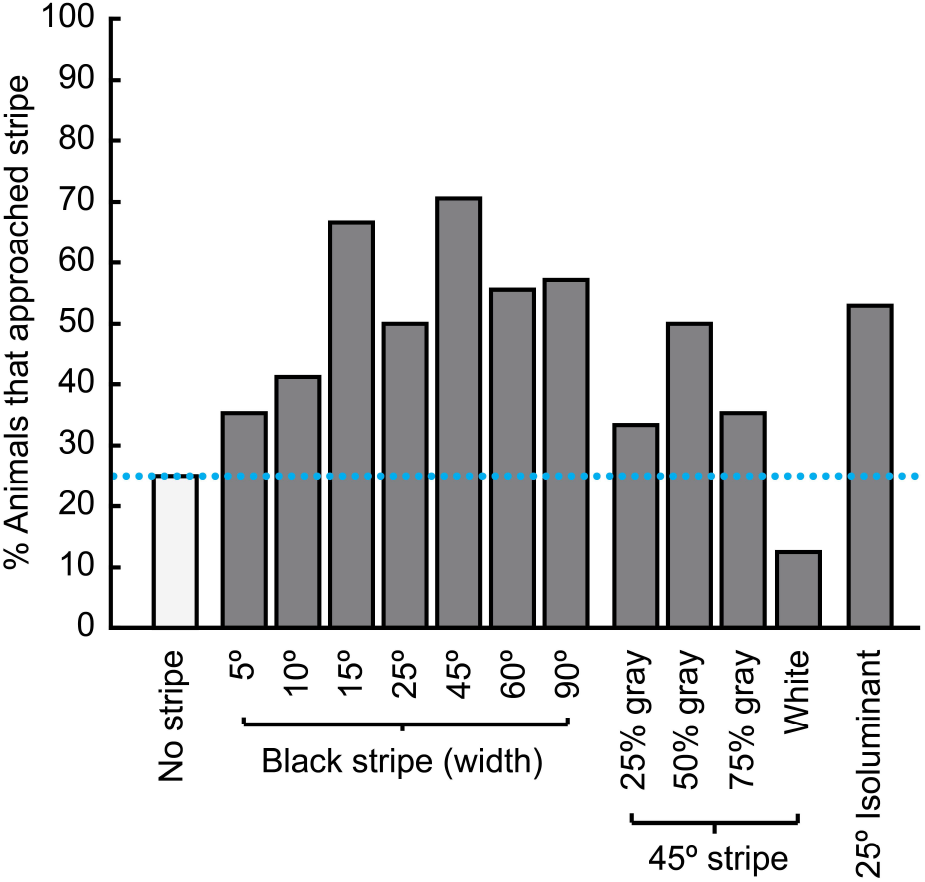
Approach rates for various visual targets. The percentage of animals that reached the wall in the quadrant centered by the stripe. When no stripe was present, the percentage of animals that reached a randomly chosen quadrant and semicircle was reported. Animals were tested with black stripes of various widths ranging from 5° to 90° (left), 45° stripes of various contrasts, and a 25° stripe that was isoluminant with the background (right). The probability of entering a random quadrant (25%) or semicircle (50%) are indicated (horizontal dashed lines). Sample sizes were *n*=17 for none; *n*=17, 17, 15, 16, 17, 18, and 14 for 5°, 10°, 15°, 25°, 45°, 60°, and 90°, respectively; *n*=12, 15, 17 for 25%, 50%, and 75% gray, respectively; *n*=18 for white; *n*=17 for isoluminant.

### *Berghia* used spatial vision

*Berghia* could be approaching visual targets through non-visual phototaxis or by using coarse spatial vision. Spatial vision is required for the detection of a visual target that is isoluminant with the background. An isoluminant visual target was created by surrounding a 25° black stripe with two 12.5° white stripes on a 50% gray background, so that the average luminance over the 50° is the same as the rest of the arena. Animals approached this isoluminant stripe near the target (Fig. 6B). A similar percentage of animals approached the 25° black stripe on an isoluminant background as approached a 25° black stripe on a white background (Fig. 5). This suggests that *Berghia* has spatial vision rather than just sensing light and dark.

**Figure 6.**
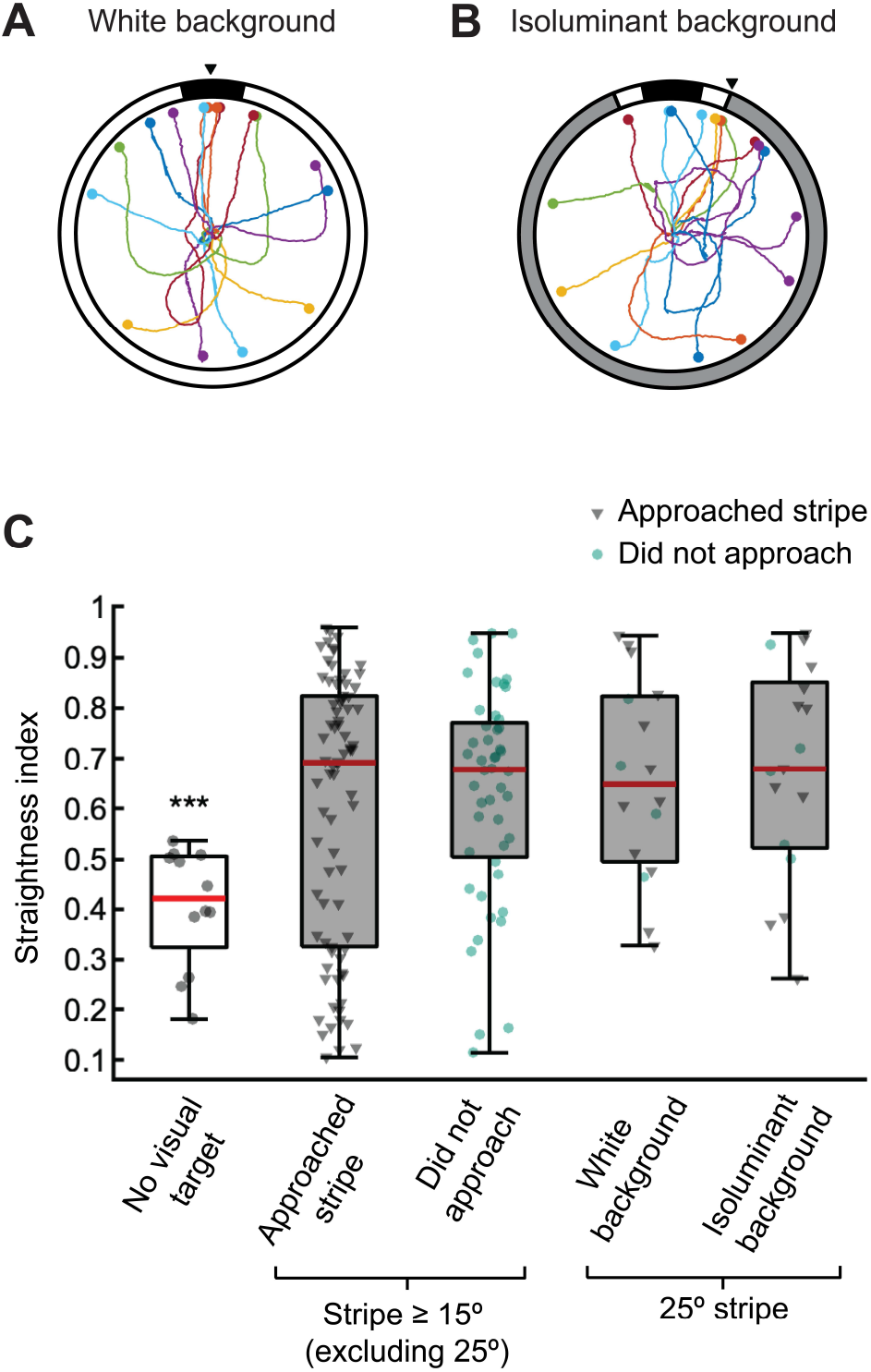
*Berghia* travelled in straighter paths with an isoluminant visual target. **A-B**. Trajectories of animals crawling from the center to the edge of an arena with a 25° black stripe on a white background (**A**) and isoluminant to the background (**B**). Maximum likelihood analysis of circular data was used; black triangles mark the direction(s) of best-fitting model for unimodal and bimodal models. Sample sizes were *n*=16 for a white background and *n*=17 for an isoluminant background. **C**. The straightness index was calculated for each animal’s path from the center to the edge of the arena. Higher values indicate a straighter path. Animals were tested without a visual target (*n*=12) or with a dark stripe on a white background of at least 15°. Animals tested with a stripe were separated by whether they approached the edge of the arena within 90° of the center of the stripe (*n*=78) or not (*n*=48). Additionally, animals were tested with a 25° black stripe on a white background (*n*=16), and a 25° stripe isoluminant to the background (*n*=17). Animals tested with a 25° stripe were marked by whether they approached (gray triangle) or did not approach the stripe (red circle) but were plotted together due to the small sample size. Box plots report the median value (red line). A one-way ANOVA (*F*=3, *P*=.02; for detailed results see Table S3-S4) with Fisher’s Least Significant Difference Test was used to compare straightness across groups; statistical differences: ***P<.001.

### *Berghia* used visual landmarks for navigation

Animals travelled in straighter paths when a visual target was present even if they did not approach it. For example, although only about half of the animals approached a 25° stripe, all of those animals travelled in a straight path to the edge of the arena (Fig. 6A). Similarly, although a stripe that was 5° did not elicit approach, several individuals travelled in direct paths to the edge (Fig. 4E). Animals also travelled in straight paths when presented with a gray stripe that was only 25% darker than the white background, which was below the contrast threshold for which animals began approaching a visual target (Fig. 4G).

The paths were straighter when a visual target of at least 15° was present (Fig. 6C). Straightness was significantly higher both for animals that approached the target and for animals that did not approach the target compared to animals tested without a visual target. Straightness was also significantly higher in animals tested with an isoluminant visual target than with no target, suggesting that animals are responding to contrast rather than luminance (Fig. 6C). Thus, *Berghia* seems to use contrasting visual landmarks as a navigational aid, even when they do not approach it.

### Visually-guided behaviors were state- and context-dependent

*Berghia’s* preference for being in the dark changed with food-deprivation and the presence of a food odor. Animals were either fed or food-deprived for 5 days and tested in normal seawater or water that was conditioned with a food odor (Fig. 7). Animals spent most of their time in the dark with food-deprivation, the presence of food odor, or both (Fig. 7C). Food-deprivation alone also seemed to strengthen the preference for dark, as 8 of 15 (53.3%) food-deprived animals spent the entire 10-minute period in the dark, whereas this occurred in only 3 of 15 (20%) fed animals. However, 4 of 15 (26.7%) animals that were food-deprived and given a food odor spent a majority of time in the illuminated side, whereas this did not occur with any of the fed animals and was observed in only 2 of 15 (13.3%) food-deprived animals and 1 of 15 (6.7%) animals tested with a food odor. This suggests that food-deprivation strengthens the preference for dark whereas the combination of being food-deprived and sensing a food odor reduces it.

**Figure 7.**
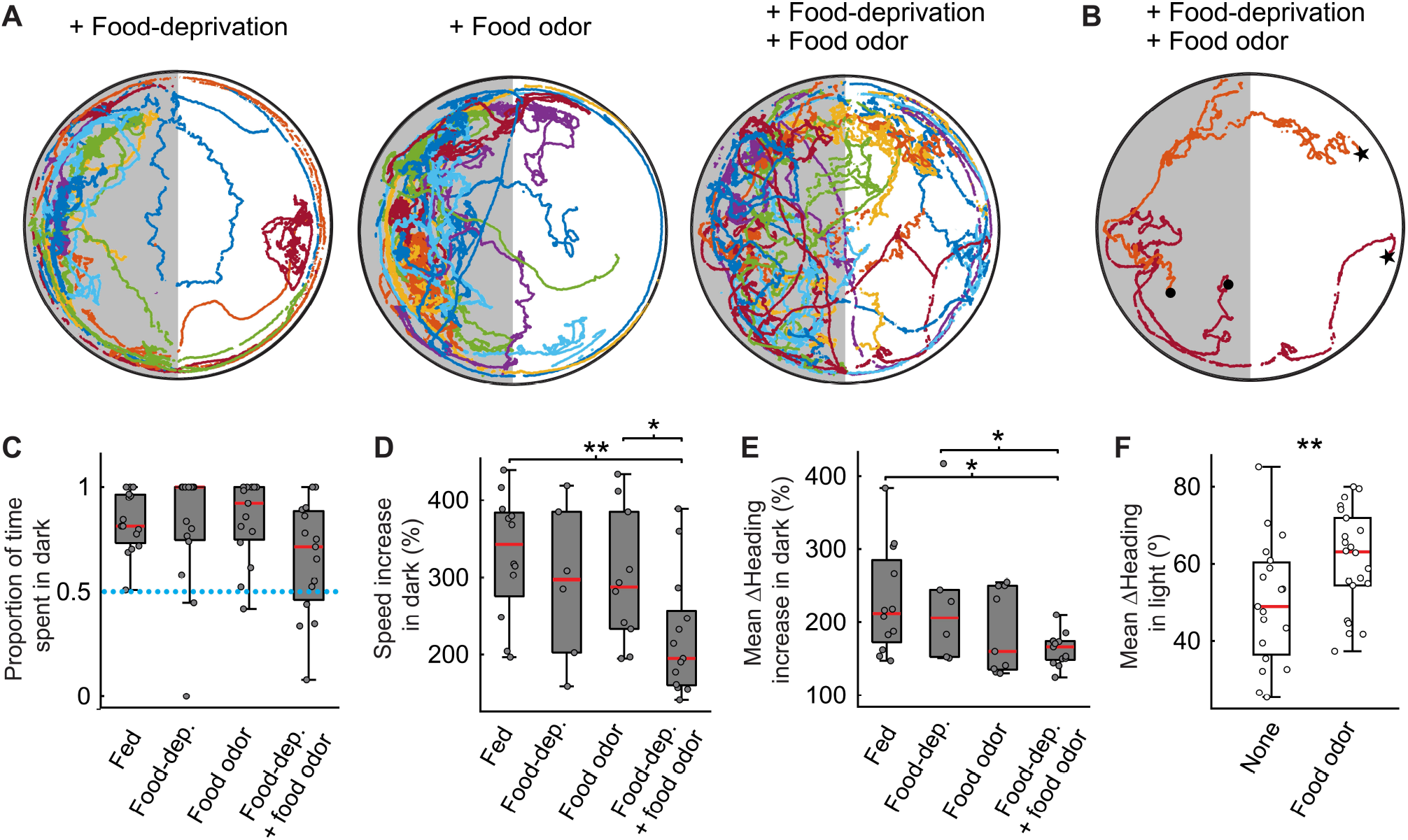
Effects of food-deprivation and food odor on behavior in a half-shaded arena. **A.** Trajectories of animals crawling in a half-shaded arena after food-deprivation (*n*=15), in the presence of food odor (*n*=15), and both of these conditions (*n*=15). **B**. Trajectories from two animals that were food-deprived and presented with a food odor. The starting position (star) and ending position (circle) of each individual is marked. **C-E**. For animals that were fed or food-deprived and tested with or without a food odor, comparisons were made of the proportion of time spent in the dark (**C**), the percentage increase in the mean speed in the dark compared to the light (**D**), and the percentage increase in the mean change in heading angle in the dark compared to the light (**E**; for separate plots of the mean speed and mean change in heading in the light and dark side, see Fig. S2). **F**. Mean change in heading angle in the light for animals tested without (*n*=30) or with a food odor present (*n*=30). A Kruskal Wallis test (**C**), two-way ANOVA with a Fisher’s Least Significant Difference Test (**D**,**E**; for detailed results see Table S5-S8), and two-sample *t*-test (**F**) were used; statistical differences: \**P*<.05, ** *P*<.01. Box plots report the median value (red line).

There were additional changes in the behavior of animals following food-deprivation and/or exposure to a food odor. All animals crawled faster in the dark than the light, regardless of feeding state or whether a food odor was present. However, this increase was significantly lower in animals that were both food-deprived and given a food odor (Fig. 7D). Additionally, whereas all animals had an increased mean change in heading angle in the dark compared to the light, this difference was smaller for food-deprived animals that were given a food odor in comparison to animals tested without a food odor (Fig. 7E). There was a significant main effect of food odor on this heading increase (Table S7). In particular, animals that were given a food odor had a larger mean change in heading in the light side of the arena compared to animals that were tested without a food odor (Fig. 7F). Thus, in addition to having a weaker preference for the dark, food-deprived animals that were given a food odor behaved more similarly in the light and dark.

Food-deprivation and sensing a food odor also reduced *Berghia’s* propensity to approach a stripe. Animals that were food-deprived approached the edge randomly, with or without a food odor (Fig. 8A). Fed animals that were given a food odor approached a 45° stripe (Fig. 8A). Just over half of the animals approached the quadrant with the stripe, while 22.2% did not approach the edge at all, suggesting that there was a reduction in the propensity to approach the stripe when animals were given a food odor (Fig. 8B). Additionally, fed animals that were given a food odor made notably sharper turns and sometimes reversed directions completely rather than travelling directly to the edge, however this was not observed in food-deprived animals that were given a food odor (Fig. 8A). Fed animals travelled in more direct paths with a stripe than without a stripe, both with and without a food odor (Fig. 8C). Food-deprived animals travelled in straighter paths than fed animals (two-sample two-tailed *t*-test, *P* = .04), but travelled with a similar straightness regardless of whether a visual target was present (Fig. 8C). Finally, fed animals crawled significantly faster with a stripe than without a stripe, however this was not true for food-deprived animals, animals given a food odor, or animals that were both food-deprived and given a food odor (Fig. 8D). Overall, animals that were food-deprived or given a food odor were less responsive to visual cues.

**Figure 8.**
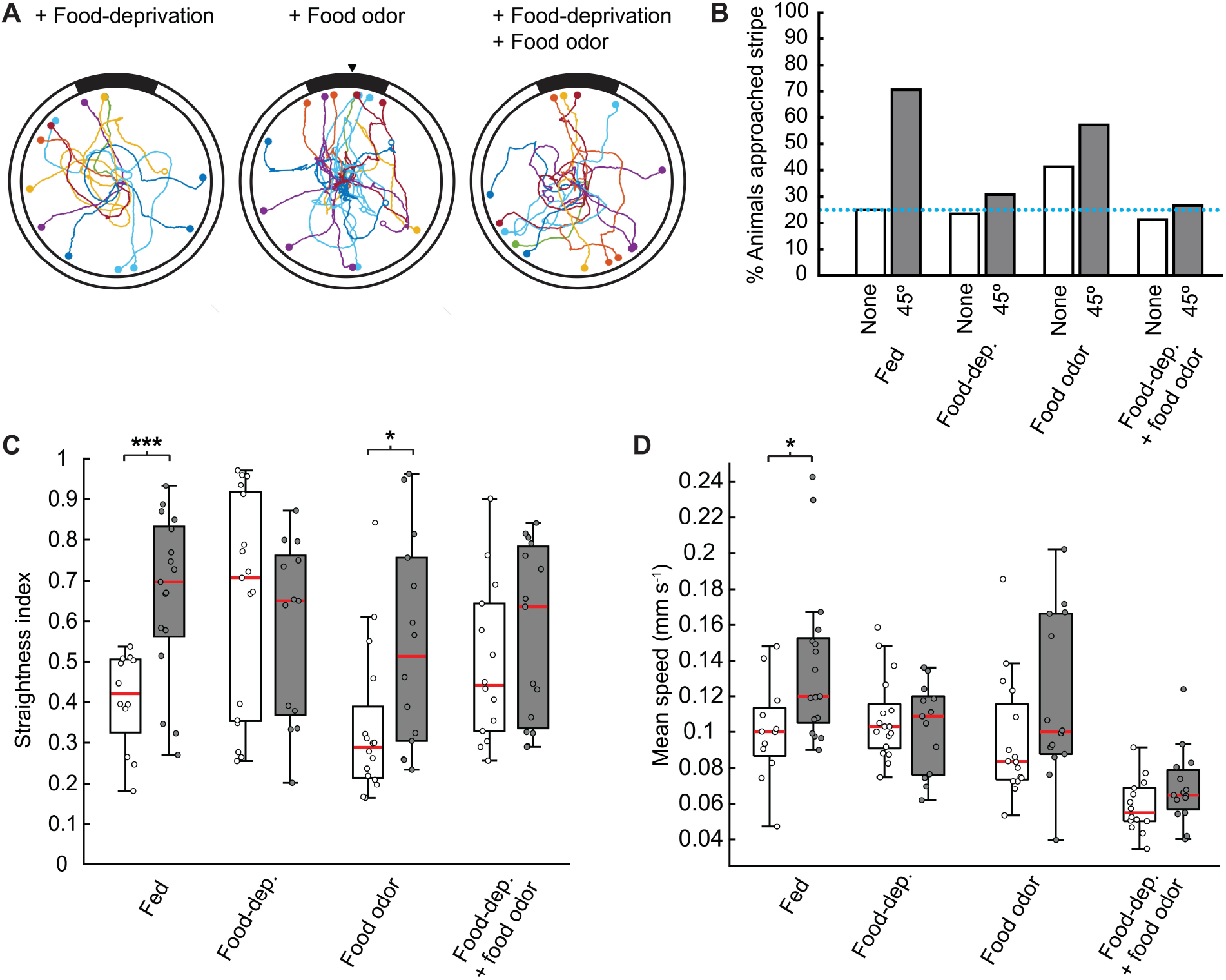
Effects of food-deprivation and food odor on the approach of visual targets. **A.** Trajectories of animals crawling in the stripe assay following food-deprivation (*n*=14), the addition of a food odor (*n*=14), or both (*n*=15). The location that each animal approached the edge of the arena is marked (colored circles). Animals that did not reach the edge within the allotted time (open circles) were plotted but excluded from analysis. Maximum likelihood analysis of circular data was used (Table S2); black triangles mark the direction(s) of best-fitting model for unimodal and bimodal models. **B.** The percentage of animals that entered the semi-circle and the quadrant containing the stripe were quantified for each combination of food-deprivation and food odor. For each group, percentages were quantified for animals tested with a visual target (diagonally lined bars) and without a visual target. When no stripe was present, the percentage of animals that entered a randomly chosen quadrant was reported. Results from fed animals with no stripe and a 45° black stripe are repeated from Figure 5. **C-D.** In addition to testing animals with a visual target, another set of animals were tested without a visual target following food-deprivation (*n*=17), the addition of a food odor (*n*=13), or both (*n*=14). For each feeding or odor condition, straightness index (**C**) and mean speed (**D**) was compared between animals tested without a visual target (white boxes) and with a visual target (gray boxes). Box plots report the median value (red line). A two-sample *t*-test (**C,D**) was used; statistical differences: \**P*<.05, \*\**P*<.01.

## Discussion

We found that *Berghia* exhibits visually-guided behaviors. Animals spent more time in dark environments and approached a contrasting visual target. When a visual target was present, animals crawled in straight paths even when they did not approach it, suggesting that visual cues are important for navigation. Animals that were food-deprived or given a food odor had a reduction in behavioral responses to visual stimuli, demonstrating that visual responses are state- and context-dependent. Additionally, there was an even stronger reduction in behavioral responses when animals were both hungry and encountered a food odor, indicating an interaction between visual information, olfactory information, and hunger state.

### Visual navigation

When given a choice between light and dark areas,*Berghia* spent most of its time in the dark and had distinctive behaviors in the light versus the dark. For example, animals followed along the edge of the arena when exposed to ambient light, which was rarely observed in the dark. Commonly referred to as thigmotaxis, this behavior is a spatial navigation strategy that has been observed in other animals, including insects (Jin et al., 2020), fish (Champagne et al., 2010; Sharma et al., 2009), rodents (Simon et al., 1994; Treit & Fundytus, 1988), and humans (Kallai et al., 2005, 2007). Thigmotaxis is thought to be performed when animals are trying to avoid or escape an environment. In *Berghia,* thigmotaxis appeared to be involved in helping animals leave illuminated environments; it was more prevalent in a partially illuminated arena, than one that was uniformly lit (Fig 3C).

*Berghia* also approached a dark vertical stripe on a light background. Although other gastropods have similarly been shown to approach a dark vertical stripe (Chiussi & Díaz, 2002; Hamilton & Winter, 1982; Hamilton & Winter, 1984; Shepeleva, 2013), this is the first demonstration of this behavior in a species of nudibranch. Similar responses in other gastropods have been suggested to be related to seeking shelter (Chiussi & Díaz, 2002) or habitat selection (Hamilton & Winter, 1982; Shepeleva, 2013). Anecdotal observations in the laboratory suggest that *Berghia* prefers to spend most of its time dark areas, such as in dark crevices and underneath objects. Additionally, *Berghia* feeds on anemones that are found in shaded areas on the roots of mangrove trees (Bedgood et al., 2020; Bellis et al., 2018). Thus, it is likely that *Berghia* approaches visual targets to seek out dark habitats that provide food and shelter.

Animals travelled in a straight path when a visual target was present even if they did not approach it, suggesting that *Berghia* uses visual landmarks to navigate its environment. External cues are indispensable in allowing animals to navigate in a straight line (Cheung et al., 2007, 2008). In addition to approaching objects, moving in a straight path allows animals to navigate to new locations, whereas convoluted paths may lead animals to re-enter previously explored areas. In the absence of directional sensory information, even humans fail to navigate in a straight path (Dacke & el Jundi, 2018). When *Berghia* was placed into an illuminated arena without any visual targets, animals changed direction several times before reaching the edge. The tortuosity of *Berghia*’s path could be a result of the arena being void of directional olfactory or visual information.

In addition to approaching objects, moving in a straight path allows animals to navigate to new locations, whereas convoluted paths may lead animals to re-enter previously explored areas. After we deprived *Berghia* of food for 5 days, animals crawled with straighter paths than animals that were regularly and recently fed. Moving in a straight path may facilitate animals travelling to new locations when food is scarce and could thus be beneficial for animals that are hungry.

### Visual capabilities of nudibranchs

In this study, we provide evidence that *Berghia* is capable of low-resolution spatial vision. Differences in *Berghia’s* behavior in the light and dark were stronger when light in the environment varied spatially than when it was uniformly illuminated. *Berghia* most effectively approached a black stripe subtending an arc of 45° around the arena while thinner or wider stripes were approached less, suggesting that *Berghia* is not simply moving toward darkness. Additionally, animals approached a 25° stripe that was isoluminant with the background, which suggests the detection of contrast rather than light intensity. It is therefore likely that *Berghia* uses spatial vision to detect objects in the environment.

Studies of the anatomy and electrophysiology of nudibranch eyes provide potential neural mechanisms that could underlie spatial vision. Although nudibranchs lack an organized retina, the microvillous regions of photoreceptors in nudibranchs form distinct areas within the eye (Dennis, 1967; Hughes, 1970; Stensaas et al., 1969). Further, photoreceptor cells in *Hermissenda* have been shown to have distinct receptive fields (Dennis, 1967). Additionally, there are inhibitory connections between the five photoreceptor cells in each eye (Crow & Tian, 2003; Detwiler & Alkon, 1973). Photoreceptor cells also inhibit cells in the contralateral optic ganglion, demonstrating a convergence of visual information between the two eyes (Alkon, 1973). It was suggested that inhibition between photoreceptors or contralateral optic ganglia may support the detection of contrast (Alkon, 1973). Thus, nudibranchs may have the necessary components for spatial vision, and the results from the current study provide behavioral evidence to support this.

### State and context dependence

Visually-guided behaviors in *Berghia* are influenced by hunger state and the sensation of food odor. When water was conditioned with *Berghia’s* prey, a sea anemone, animals still preferred dark environments, but they had a slight reduction in their propensity to approach a black stripe. Additionally, they showed changes in the style of locomotion, with animals performing sharper rather than smooth turns following the addition of a food odor. Following food-deprivation, animals did not approach a black stripe. When both food-deprived and given a food odor, animals had a weaker preference for being in the dark and behaved more similarly in the light and dark. Additionally, fed animals that were tested with a food odor often sharply reversed directions, while this never occurred in food-deprived animals that were given a food odor. Together, these results indicate that there are interactions between hunger state, olfactory information, and visual information that lead to changes in *Berghia*’s behavior.

Visual responses in other gastropods are also dependent on internal state or the presence of olfactory information. Similar to *Berghia,* the sea snail *Nerita fulgarans* approaches dark visual targets. When presented with a predator odor, *Nerita* avoids rather than approaches visual stimuli (Chiussi & Díaz, 2002). The dorid nudibranch *Chromodoris zebra* ceased orienting to light when they were with conspecifics (Crozier & Arey, 1919). The sea slug *Pleurobranchaea californica* responds to food preferentially to light (Davis & Mpitsos, 1971). The aeolid nudibranch *Hermissenda crassicornis* changes its preference for light according to the time of day, with bright light being preferred during the day but not during the night (Lederhendler et al., 1980). When given a choice, *Hermissenda* approaches a light over a food source, except when hungry (Alkon et al., 1978). Furthermore, stimulation of tentacular chemoreceptors inhibits the responses of photoreceptor cells in *Hermissenda* (Alkon et al., 1978). In these gastropods, as well as *Berghia*, visual information seems to be ignored during other behaviors such as foraging or seeking mates.

### Conclusion

Although vision was not previously considered an important sensory modality for nudibranchs, the current study provides behavioral evidence that nudibranchs respond to visual features of their environment. Our findings demonstrate that *Berghia* has visually-guided behaviors that are influenced by hunger state and odors. It is likely that *Berghia* uses its eyes for low-resolution visual tasks such as seeking dark habitats, approaching objects, and navigating its environment.

## Acknowledgments

We thank Amanda Cho for assistance with designing the half-shaded arena and collecting pilot data. We also thank Niah Holtz and Jackson Southard for assistance with data collection. Additionally, we thank Thi Bui for determining *Berghia’s* sensitivity to long-wavelength light.

## Data availability

Data and analysis code available upon request.

